# Bacterium secretes chemical inhibitor that sensitizes competitor to bacteriophage infection

**DOI:** 10.1101/2024.01.31.578241

**Authors:** Zhiyu Zang, Chengqian Zhang, Kyoung Jin Park, Daniel A. Schwartz, Ram Podicheti, Jay T. Lennon, Joseph P. Gerdt

## Abstract

To overtake competitors, microbes produce and secrete secondary metabolites that kill neighboring cells and sequester nutrients. This natural product-mediated competition likely evolved in complex microbial communities that included viral pathogens. From this ecological context, we hypothesized that microbes secrete metabolites that “weaponize” natural pathogens (i.e., bacteriophages) to lyse their competitors. Indeed, we discovered a bacterial secondary metabolite that sensitizes other bacteria to phage infection. We found that this metabolite provides the producer (a *Streptomyces* sp.) with a fitness advantage over its competitor (*Bacillus subtilis*) by promoting phage infection. The phage-promoting metabolite, coelichelin, sensitized *B. subtilis* to a wide panel of lytic phages, and it did so by preventing the early stages of sporulation through iron sequestration. Beyond coelichelin, other natural products may provide phage-mediated competitive advantages to their producers—either by inhibiting sporulation or through yet-unknown mechanisms.

## Introduction

Competition is a common theme in microbial life.^1^ Microbes frequently live in complex communities with diverse bacterial, fungal, and protozoan species.^2^ Given the finite resources and space in their niches, microbes have evolved an arsenal of strategies that allow them to persist in the face of competition.^3^ The secretion of secondary metabolites is one of the most common methods by which bacteria and fungi compete with neighboring microbes.^1, 3^ For example, antibiotics can kill or arrest the growth of competitors.^4^ Anti-adhesion molecules like biosurfactants can impede invasion and colonization from potential competitors.^5^ Siderophores that harvest the limited iron in the environment can also indirectly starve competitors and prevent their rapid growth.^6^

To find new mechanisms of natural product-based microbial competition, we considered environmental factors that microbes might leverage for a competitive advantage. Since microbes naturally compete within complex populations that involve interactions with other predators and pathogens, we asked if microbes secrete metabolites that sensitize their competitors to the ubiquitous predators or pathogens around them. The major pathogens of bacteria are bacteriophage viruses.^7^ These obligate parasites are strong agents of selection that can induce high rates of mortality and have indirect effects on the competition between microbes and the flux of resources in their environment.^8^ Bacteria have evolved myriad defenses that confer immunity to virus infection.^9^ We hypothesize that microbes may inhibit these antiviral defenses of their competitors, thereby sensitizing their competitors to phages to gain a relative fitness advantage. A similar form of “weaponizing” phages has been reported in which the secretion of secondary metabolites by one species induces lysogenic phages to become lytic and kill the host of another species.^10–12^ Otherwise, most cases report that secondary metabolites protect bacteria from phage infection (e.g., anthracyclines and aminoglycosides have exhibited antiviral effects).^13^ To identify cases where a microbe sensitizes competing bacteria to phages, we screened soil bacteria for isolates that improved the infectivity of bacteriophages on the model soil bacterium *Bacillus subtilis*.

We discovered that a *Streptomyces* sp. outcompetes *B. subtilis* in laboratory culture by secreting a metabolite that sensitizes *B. subtilis* to phage predation. The metabolite is a common siderophore named coelichelin, which elicited its effect via iron sequestration. We further found that the improved phage predation was due to the ability of coelichelin to delay sporulation of *B. subtilis*, which broadly sensitized *B. subtilis* to many lytic phages. This finding supports the hypothesis that sporulation is a phage-defense strategy.^14–16^ Furthermore, given the abundance of phages and iron-sequestering metabolites, this mechanism of metabolite-induced phage sensitization may be a common approach to outcompete spore-forming bacteria.

## Results

### Metabolites of *Streptomyces sp.* promote phage plaquing in *B. subtilis*

We performed a binary-interaction screen to discover bacteria that promote phage infections in *B. subtilis*. Colonies of soil-isolated Actinobacteria were pre-grown on an agar plate to secrete metabolites into their medium (Fig. 1a). Subsequently, *B. subtilis* and phage SPO1 were plated around the mature Actinobacteria colonies. Without an adjacent Actinobacteria colony, the SPO1 plaque sizes were small, but we hypothesized that some adjacent colonies could sensitize the *B. subtilis* to the phage and enlarge its plaques. In a small screen of 54 soil-derived Actinobacteria, we found several that caused SPO1 to form larger plaques on *B. subtilis.* One of the clearest plaque-enlarging isolates was *Streptomyces* sp. I8-5 (Fig. 1b,c). Notably, the plaque sizes were most enlarged near the *Streptomyces* colony, and distant plaques were essentially the normal small size (Fig. 1c). This distance dependence suggested the production of a diffusible plaque-promoting substance—or possibly the depletion of a diffusible plaque inhibitory substance from the media. To investigate whether the promoted plaquing was due to metabolites secreted from the I8-5 colony, we tested the activity of sterile-filtered conditioned medium of I8-5 culture. Concentrated conditioned medium of I8-5 was spotted onto an agar plate containing *B. subtilis* and phage SPO1. The conditioned medium increased the plaque sizes ∼30-fold (Fig. 1d), suggesting that secreted component(s) from I8-5 promote *Bacillus* phage infection.

**Figure 1.**
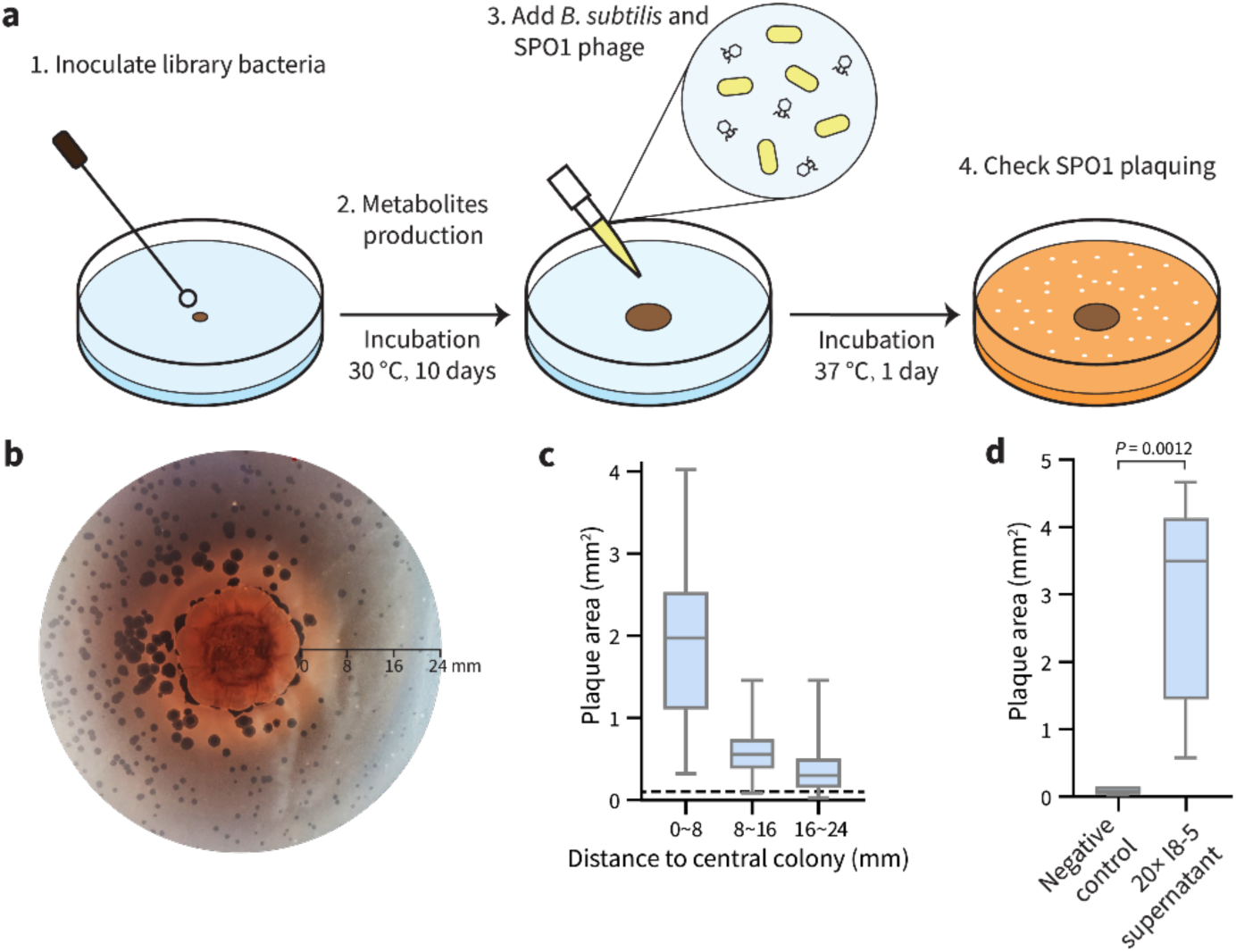
Binary-interaction screen identifies a *Streptomyces* sp. that promotes SPO1 phage predation of *B. subtilis*. (a) Scheme of the binary-interaction screen. **(b)** A mature colony of *Streptomyces* sp. I8-5 (center) promoted SPO1 phage proliferation nearby (dark circles are plaques), especially within a radius of 8 mm. **(c)** Quantification of plaque areas with increasing distance from the *Streptomyces* sp. I8-5 colony. Data are represented as boxplots, showing the median, interquartile ranges and minimum (bottom bar) and maximum (top bar). The dashed line represents the average plaque area of SPO1 phages in the absence of the *Streptomyces* colony. At least 72 plaques were measured for each condition. **(d)** The I8-5 supernatant was concentrated 20 times and tested for the ability to enlarge SPO1 plaques. Water was used as a negative control. Data are represented as boxplots, showing the median, interquartile ranges and minimum (bottom bar) and maximum (top bar). At least nine plaques were measured for each condition.

### Plaque-promoting metabolite is the siderophore coelichelin

To identify the plaque-enlarging metabolite(s) made by *Streptomyces* sp. I8-5, the conditioned medium of I8-5 culture was subjected to bioactivity-guided fractionation. Two active semi-pure fractions were obtained from different purification strategies: one from reversed-phase chromatography and the other from cation exchange chromatography (Fig. 2a). Since these are relatively orthogonal separation methods, we suspected that few metabolites would be shared between the active fractions. Therefore, we compared the composition of the two active fractions using liquid chromatography-mass spectrometry (LC-MS), and we found that only two putative metabolites were shared by the two fractions (Fig. 2a). High-resolution mass spectrometry of these two metabolites revealed one with *m/z* 566.2867 and one with *m/z* 619.1946 (both were likely [M+H]^+^ adducts due to matching [M−H]^–^ adducts observed by negative mode analysis [Extended Data Fig. 1]). Tandem mass spectrometry (MS/MS) analysis of the 566.2867 peak (Fig. 2b) revealed a fragmentation pattern that matched the known metabolite coelichelin (Fig. 2c,d).^17^ The exact mass of the parent ion also matched the expected [M+H]^+^ mass of coelichelin within 15 ppm error. Because coelichelin is a siderophore with high affinity to iron,^17, 18^ the *m/z* 619.1946 peak was attributed to the Fe-coelichelin complex [M−2H+Fe]^+^ with 8 ppm error. To confirm the ability of *Streptomyces* sp. I8-5 to produce coelichelin, we sequenced its genome and identified the coelichelin biosynthetic gene cluster (BGC) (Fig. 2e). The coelichelin BGC in I8-5 has the same organization as the reported one with high sequence identity (>75% for each gene, Fig. 2e).^17^ Consistent with the reported coelichelin non-ribosomal peptide synthetase (NRPS),^17^ epimerization domains were identified in the first and second module of the I8-5 coelichelin NRPS (Fig. 2e). These domains strongly suggest that the absolute stereochemistry of the four amino acids in our sample are the same as reported (Fig. 2c).^17^

**Figure 2.**
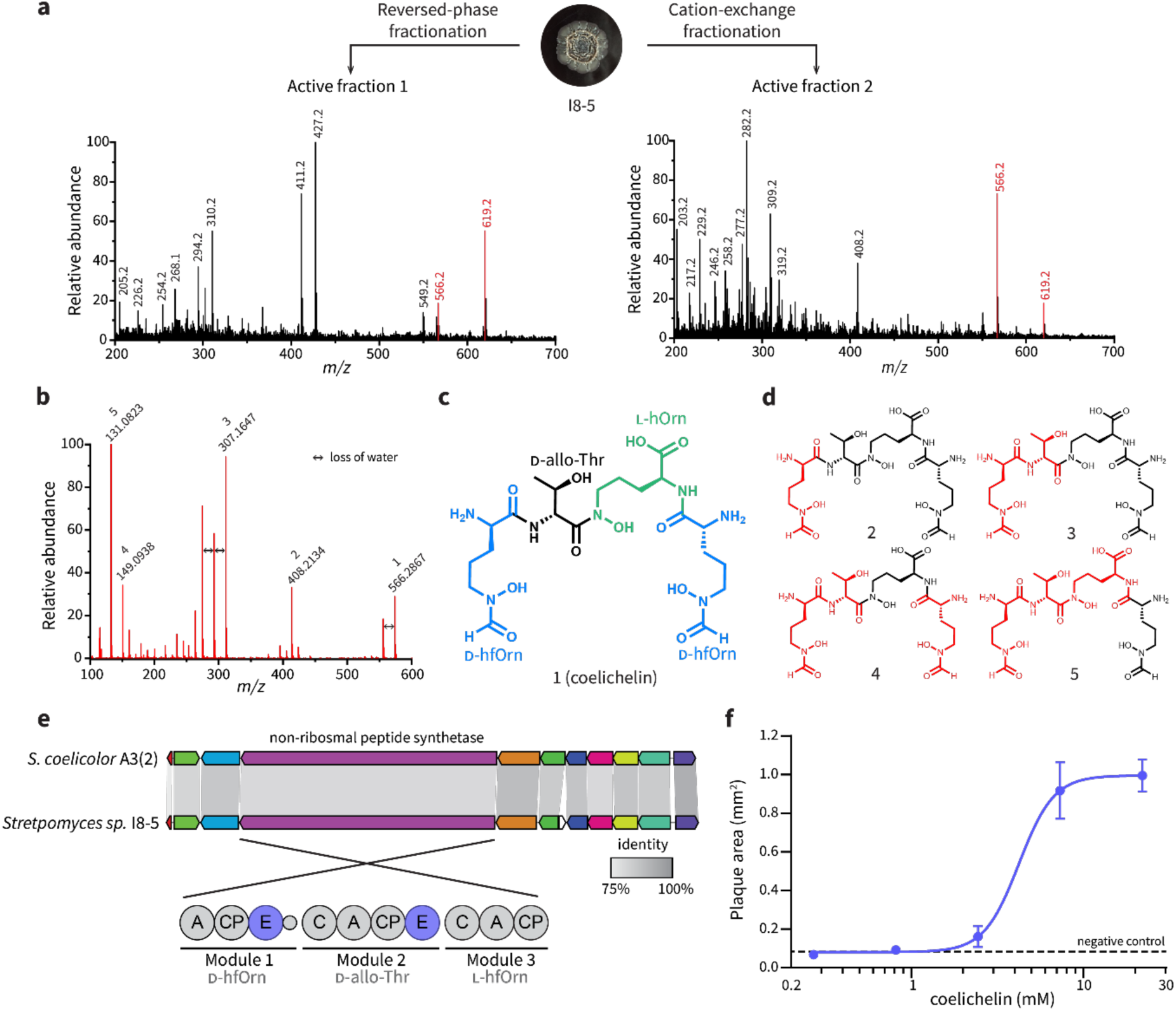
Coelichelin is the active metabolite that promotes phage predation. **(a)** Bioactivity-guided fractionation and MS analysis identified only two putative metabolites present in both active fractions purified from orthogonal separation techniques. Positive mode electrospray ionization results are shown here, and matching negative mode peaks (*m/z* 564.2 and 617.2) are shown in Extended Data Fig. 1. **(b)** MS/MS spectrum of the *m/z* 566.2867 peak, matching that of the [M+H]^+^ adduct of coelichelin.^17^ **(c)** Chemical structure of coelichelin, highlighting each amino acid residue. **(d)** Key fragment losses of coelichelin in the MS/MS analysis, annotated with their associated peak number from panel (b). Black atoms indicate the observed fragment ions, and the neutral lost fragments are highlighted in red. **(e)** Comparison of the *Streptomyces* sp. I8-5 coelichelin biosynthetic gene cluster with the reported one from *S. coelicolor* A3(2). The percent identity between each pair of genes is shown with shading (all were >75%). The modules of the coelichelin non-ribosomal peptide synthetase are shown in detail below. The three modules are responsible for installation of D-δ-*N*-formyl-δ-*N*-hydroxyornithine (D-hfOrn), D-allo-threonine (D-allo-Thr), and L-δ-*N*-hydroxyornithine (L-hOrn), respectively. The adenylation domains (A), thiolation and peptide carrier proteins (CP), condensation domains (C), and epimerization domains (E) are shown. **(f)** Pure coelichelin enlarged phage plaques in a dose-dependent manner (EC_50_=4.2 mM). Water was used as a negative control. Data are represented as the average ± SEM from at least seven individual plaques of each condition.

To determine if coelichelin was the active component secreted by *Streptomyces* sp. I8-5, we purified coelichelin from the *Streptomyces*-conditioned medium (Extended Data Fig. 2). NMR analyses were performed on the gallium complex of the purified coelichelin to afford clean spectroscopic data. These analyses confirmed the identity and purity of our isolated coelichelin (Extended Data Fig. 3, Extended Data Table 1, and Supplementary Figures 1–4). As hypothesized, the pure coelichelin enlarged plaques in a dose-dependent manner, confirming it is a phage-promoting metabolite secreted by *Streptomyces* sp. I8-5 (Fig. 2f).

### Coelichelin promotes phage proliferation by iron sequestration

Coelichelin is known to be a hydroxamate-type siderophore.^17–20^ We hypothesized that other siderophores would also enlarge plaques of SPO1. We tested three other common siderophores: ferrichrome, enterobactin, and linear enterobactin (Fig. 3a). Surprisingly, none of these three siderophores promoted phage plaquing (Fig. 3b). Previous work has demonstrated that *B. subtilis* can import a range of xenosiderophores produced by other organisms (in addition to using its own siderophore bacillibactin).^21^ This list of “pirated” siderophores notably includes all three that failed to enlarge plaques: ferrichrome,^22^ enterobactin,^23, 24^ and linear enterobactin.^22^ Therefore we hypothesized that only siderophores that cannot be imported and utilized by *B. subtilis* enlarge plaques. The non-usable siderophores would sequester iron away from *B. subtilis* and, via an unknown mechanism, improve phage replication on the iron-starved host.

**Figure 3.**
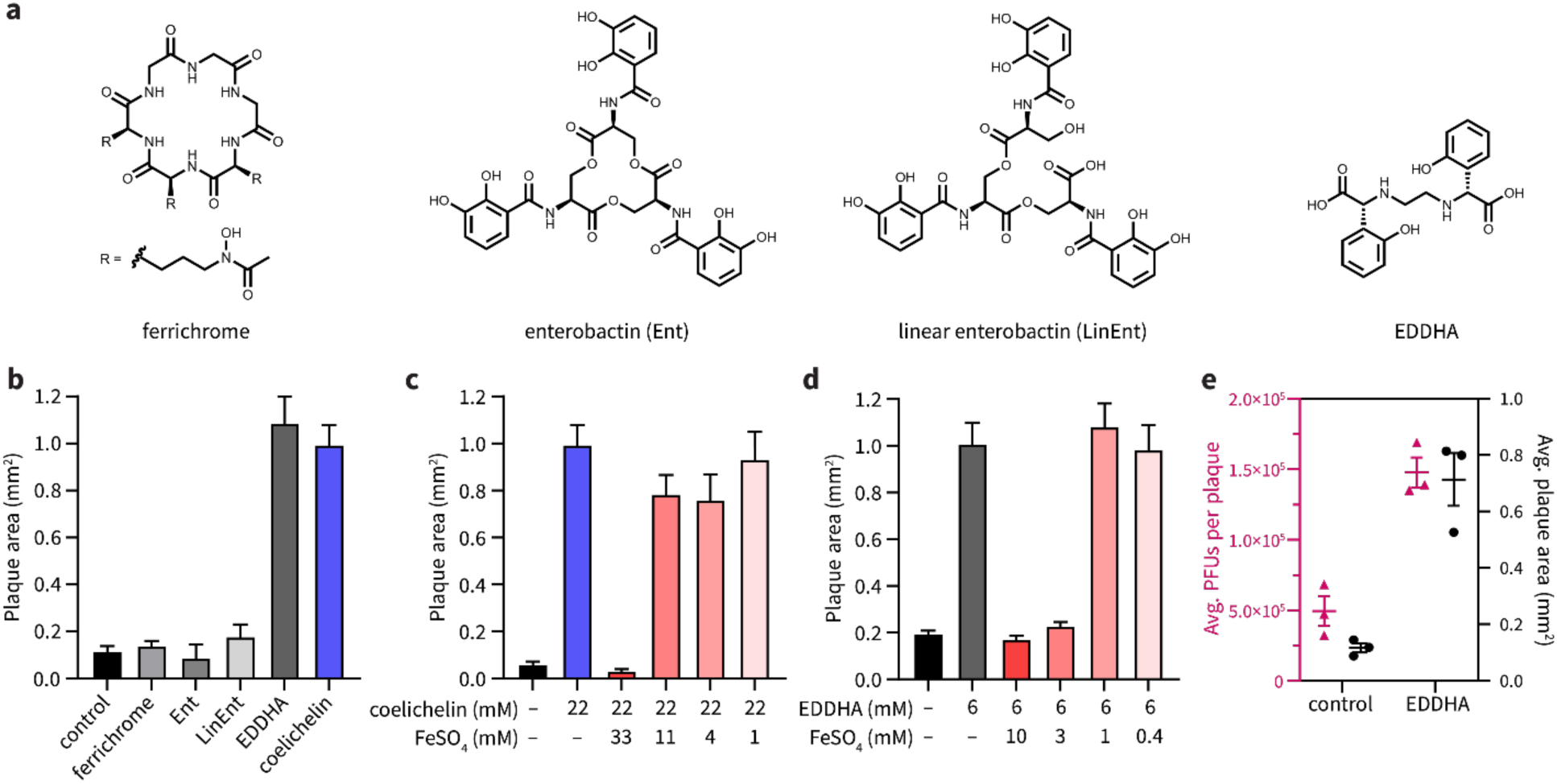
Coelichelin promotes phage predation by sequestering iron. **(a)** Chemical structures of ferrichrome, enterobactin (Ent), linear enterobactin (LinEnt), and ethylenediamine-*N,N*′-bis(2-hydroxyphenyl-acetic acid) (EDDHA). **(b)** Ferrichrome (20 mM), Ent (10 mM), LinEnt (20 mM), and EDDHA (6 mM) were tested for the ability to increase SPO1 plaque areas. Water was used as a negative control. Data are represented as the average ± SEM from at least four individual plaques of each condition. **(c)** Iron complementation antagonized the plaquing promotion effect of coelichelin and **(d)** EDDHA. Water was used as a negative control. Data are represented as the average ± SEM from at least six individual plaques of each condition. **(e)** The average plaque forming units (PFUs) per plaque and plaque area were measured with EDDHA (6 mM) or water (control) treatment. At least 13 plaques were selected for each condition. Data are represented as the average ± SEM from three independent biological replicates. Symbols show the values of each biological replicate.

It was previously unknown if coelichelin could sequester iron away from *B. subtilis* (or if *B. subtilis* could instead use it as a xenosiderophore). Therefore, to test the iron sequestration hypothesis, we utilized a synthetic iron chelator, ethylenediamine-*N,N*′-bis(2-hydroxyphenyl-acetic acid) (EDDHA, Fig. 3a), which is known to sequester iron away from *B. subtilis.*^25^ In line with our hypothesis, EDDHA increased SPO1 plaque sizes to a similar level as coelichelin (Fig. 3b). Furthermore, if iron starvation is responsible for the improved plaquing, the plaque sizes should decrease to their normal size when excess iron is co-administered with the siderophore. As we expected, approximately equimolar concentrations of FeSO_4_ quenched the plaque promotion effect of both coelichelin and EDDHA. (Fig. 3c,d). We hypothesized that the enlarged plaques were the result of increased phage proliferation within each plaque. To test this hypothesis, we quantified the viable phages (plaque forming units [PFUs]) generated per plaque. Indeed, the larger plaques afforded by iron limitation produced far more viable phages (Fig. 3e). Thus, we concluded that *B. subtilis* does not use coelichelin as a xenosiderophore, and furthermore, the iron starvation caused by this *Streptomyces* siderophore promotes the predation of *B. subtilis* by the SPO1 phage.

### Iron sequestration improves phage infection by inhibiting sporulation in *B. subtilis*

We next investigated the mechanism by which iron starvation promoted phage infection of *B. subtilis.* Plaque size can be increased by many factors that either accelerate the rate of phage replication or extend the period in which phages can replicate before the bacteria becomes recalcitrant. For example, the rate of plaque expansion depends largely on the burst size (i.e., the number of new phages released from each infected cell) and latent period (i.e., the time required for phages to lyse the host cell and produce new progeny) of phage replication. Specifically, large burst sizes and shorter latent periods maximize the phage reproduction rate, thus resulting in larger plaques.^26^ We considered whether iron starvation increased burst size and/or shortened the latent period of phage replication. It has been reported that iron starvation actually has the opposite effect in *Vibrio cholerae*: it reduces burst size and delays phage-mediated cell lysis.^27^ In line with the *V. cholerae* study, we observed that when *B. subtilis* grew next to *Streptomyces* sp. I8-5, the plaque development of SPO1 was slower than plaque development alone, suggesting that iron sequestration does not accelerate phage replication (Fig. 4a). However, we observed that the plaque development process lasted longer in the presence of the *Streptomyces* colony (Fig. 4a), which means that phages underwent more reproduction cycles, lysing more *B. subtilis* cells and ultimately forming 10× larger plaque areas.

**Figure 4.**
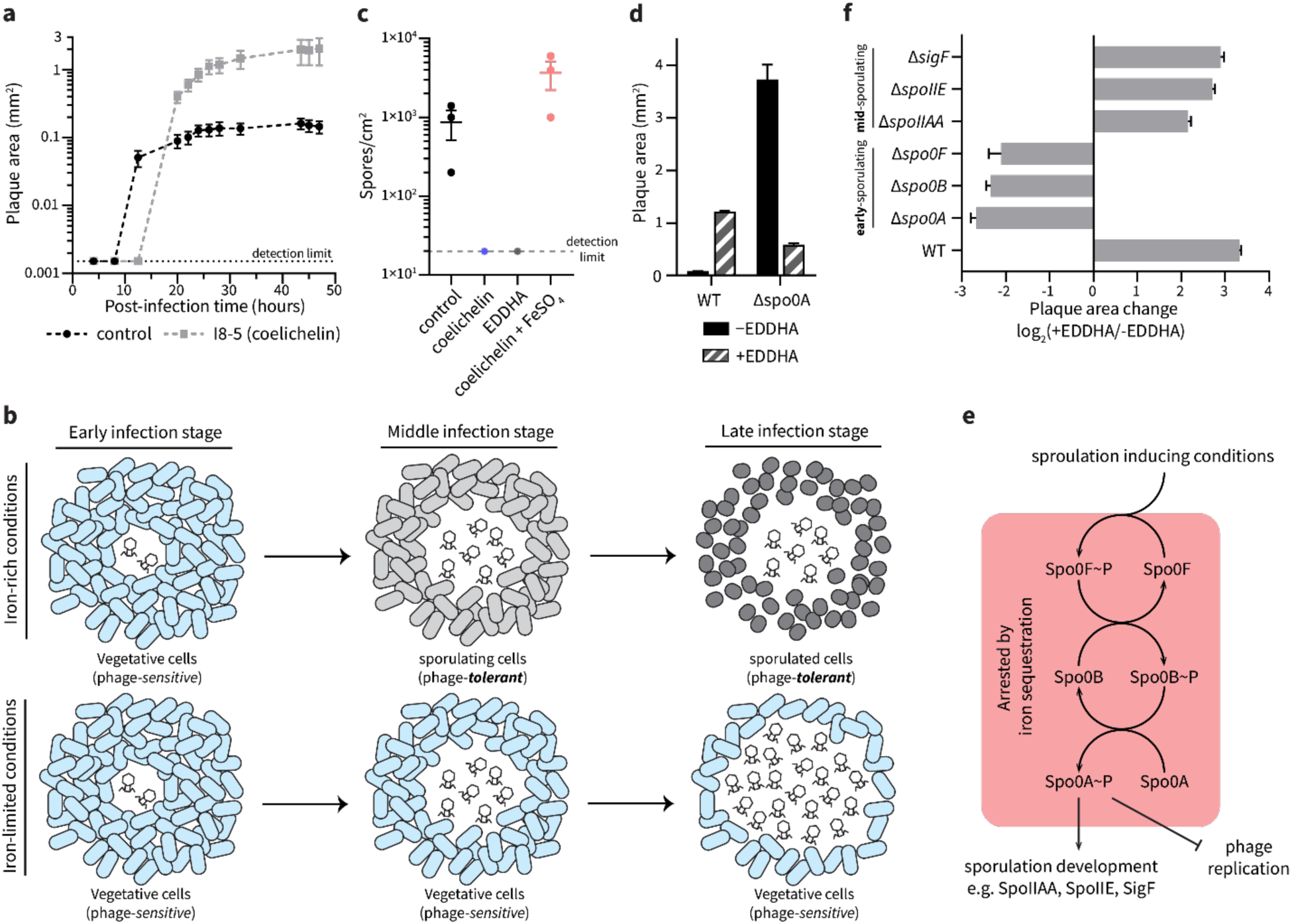
Iron sequestration inhibits sporulation initiation in *B. subtilis.* **(a)** The plaque size development of SPO1 phage on *B. subtilis* grown ∼8 mm from to an I8-5 colony (coelichelin produced) or *B. subtilis* alone (control). Data are represented as the average ± SEM from at least eight individual plaques of each condition. **(b)** Schematic model for iron sequestration-induced promotion of phage infection: under iron-rich conditions, *B. subtilis* cells sporulate when nutrients are limited. Once the sporulation process initiates in *B. subtilis* cells, phage proliferation is arrested (top). However, when iron is limited, *B. subtilis* cells are unable to sporulate, allowing phages to continue infecting vegetative *B. subtilis* cells (bottom). **(c)** The influence of iron starvation on *B. subtilis* sporulation. The number of spores formed under treatment of water (control), coelichelin (22 mM), EDDHA (6 mM), and coelichelin (22 mM) + FeSO_4_ (33 mM). Iron starvation inhibited sporulation. Data are represented as the average ± SEM from three independent biological replicates. Circles show the values of each biological replicate. **(d)** The plaque-enlarging effect of EDDHA (6 mM) was tested against *B. subtilis* WT and Δ*spo0A*. Water was used as the −EDDHA control. The Δ*spo0A* mutant naturally formed larger plaques that were not further increased by iron sequestration. Data are represented as the average ± SEM from at least six individual plaques of each condition. **(e)** Schematic representation of sporulation steps in *B. subtilis*. **(f)** The plaque size ratio between EDDHA-treated and untreated samples of different mutants. Mutations in *spo0A* and earlier genes eliminated the phage-promoting effects of iron sequestration. Data are represented as the average ratio ± SEM calculated from at least four individual plaques of each condition.

The extended period of phage infection led us to consider an alternative mechanism: iron starvation may prevent host dormancy. Phage reproduction and plaque development requires metabolically active host cells.^28^ Under sub-optimal conditions, many bacteria enter dormant stages with heavily reduced metabolic activity,^29^ which is unfavorable for phage proliferation. Therefore, dormancy can be considered a mechanism of phage defense. One form of dormancy in *B. subtilis* is sporulation,^30^ which has been shown to arrest phage infection by masking surface receptors^14^ and inhibiting DNA replication and transcription.^15, 16^ Notably, sporulation in *B. subtilis* relies on sufficient levels of intracellular iron.^31^ Therefore, we hypothesized that iron starvation inhibits the sporulation process and maintains *B. subtilis* cells in their phage-susceptible vegetative growth state. Consequently, delayed sporulation would promote phage proliferation, leading to larger plaques (Fig. 4b). To test this hypothesis, we first determined if iron sequestration inhibited sporulation of *B. subtilis* under our experimental conditions. Indeed, by quantifying heat-treated spores,^32^ we determined that both coelichelin and EDDHA inhibited sporulation in *B. subtilis* (Fig. 4c). This inhibitory effect was due to iron sequestration, as demonstrated by the ability of iron supplementation to recover native levels of sporulation (Fig. 4c).

To determine if sporulation inhibition was the cause of increased phage proliferation, we employed a knockout mutant of *spo0A* in *B. subtilis*. Since Spo0A is a transcriptional regulator that initiates the sporulation process in *B. subtilis*, Δ*spo0A* mutants are incapable of sporulating and are “locked” into their vegetative growth state.^33^ We predicted that this mutant would naturally form large plaques that are not further enlarged by iron limitation. Indeed, the SPO1 phage formed extremely large plaques on the Δ*spo0A* mutant (Fig. 4d). As expected, the mutant plaques were not enlarged by EDDHA-induced iron sequestration. In fact, the plaques were substantially smaller under EDDHA treatment, possibly due to the aforementioned slowing of phage proliferation under iron limitation.^27^ Therefore, our results demonstrate that iron sequestration extends phage infection on *B. subtilis* by inhibiting its sporulation into dormant, phage-tolerant cells.

### Iron sequestration inhibits sporulation at an early stage

Sporulation in *B. subtilis* is a multi-stage process involving initiation, development, and maturation.^34^ The process is transcriptionally initiated by Spo0A, which is activated by the kinase Spo0B, which itself is activated by the kinase Spo0F in a phosphorelay (Fig. 4e).^35^ The activated Spo0A protein then functions as the master transcriptional regulator of the genes leading to the formation of mature *B. subtilis* spores. As further confirmation of the necessity of activated Spo0A protein for the small plaque phenotype in untreated *B. subtilis*, we tested Δ*spo0F* and Δ*spo0B* mutants. These, too, exhibited large plaques that did not enlarge upon iron sequestration (Fig. 4f). Therefore, the phage inhibition effect occurs downstream of Spo0A activation, and iron sequestration likely prevents the activation of Spo0A.

To determine if the iron-dependent phage inhibition phenotype requires the complete maturation of spores, we tested mutants of three genes that are required for intermediate stages of sporulation: *spoIIAA*, *spoIIE*, and *sigF*.^36^ Unlike the early sporulation genes, these mutants plaqued like wild-type: they formed small plaques under resource-replete conditions but large plaques under iron starvation (Fig. 4f). Therefore, we propose that phage proliferation is arrested by the earliest stages of sporulation and does not require the intermediate and final stages of spore formation. Notably, prior work has shown that activated Spo0A can directly inhibit replication of and transcription from phage phi29 DNA.^15, 16^ Therefore, two likely mechanisms may contribute to the Spo0A-dependent plaque restriction: Spo0A may directly inhibit replication and transcription of phage SPO1 genes, and/or Spo0A may activate early sporulation processes that halt host metabolic processes that are essential for phage replication. These mechanisms are not mutually exclusive and may synergize to fully arrest phage replication in our conditions. In either case, iron sequestration prevents the activation of Spo0A, which frees the phage to continue replicating and lysing the *B. subtilis* population.

### Siderophore production enables *Streptomyces* to outcompete *B. subtilis* in a phage-dependent manner

*B. subtilis* and *Streptomyces* are both soil bacteria and are likely to share habitats in nature.^37^ Since coelichelin secreted by *Streptomyces* sp. I8-5 promoted phage infection on *B. subtilis* (Fig. 5a), we hypothesized that coelichelin offers *Streptomyces* a competitive advantage over *B. subtilis* in the presence of *Bacillus* phages. Indeed, the combined action of SPO1 phages and a nearby *Streptomyces* sp. I8-5 colony significantly decreased the *B. subtilis* population density relative to phage treatment or *Streptomyces* treatment alone (Fig. 5b). Importantly, this effect allowed *Streptomyces* to outcompete *B. subtilis* 15:1 under our growth conditions (Fig. 5c). To validate the significance of coelichelin-induced iron sequestration for the decreased *B. subtilis* fitness, we supplied the *B. subtilis* cells with excess bioavailable iron in the form of a xenosiderophore-iron complex with ferrioxamine E (Fig. 5d). With ferrioxamine E as a supplemented iron source, the phage promotion effect from *Streptomyces* was abolished as seen with normal size plaques (Fig. 5a), and the *B. subtilis* population increased to the same level as the no-*Streptomyces* and no-phage controls (Fig. 5b). With the lost impact of coelichelin, *Streptomyces* lost its *phage-*induced competitive advantage over *B. subtilis* (Fig. 5c). Therefore, secretion of coelichelin enables *Streptomyces* sp. I8-5 to outcompete *B. subtilis* by facilitating phage predation on its competitor.

**Figure 5.**
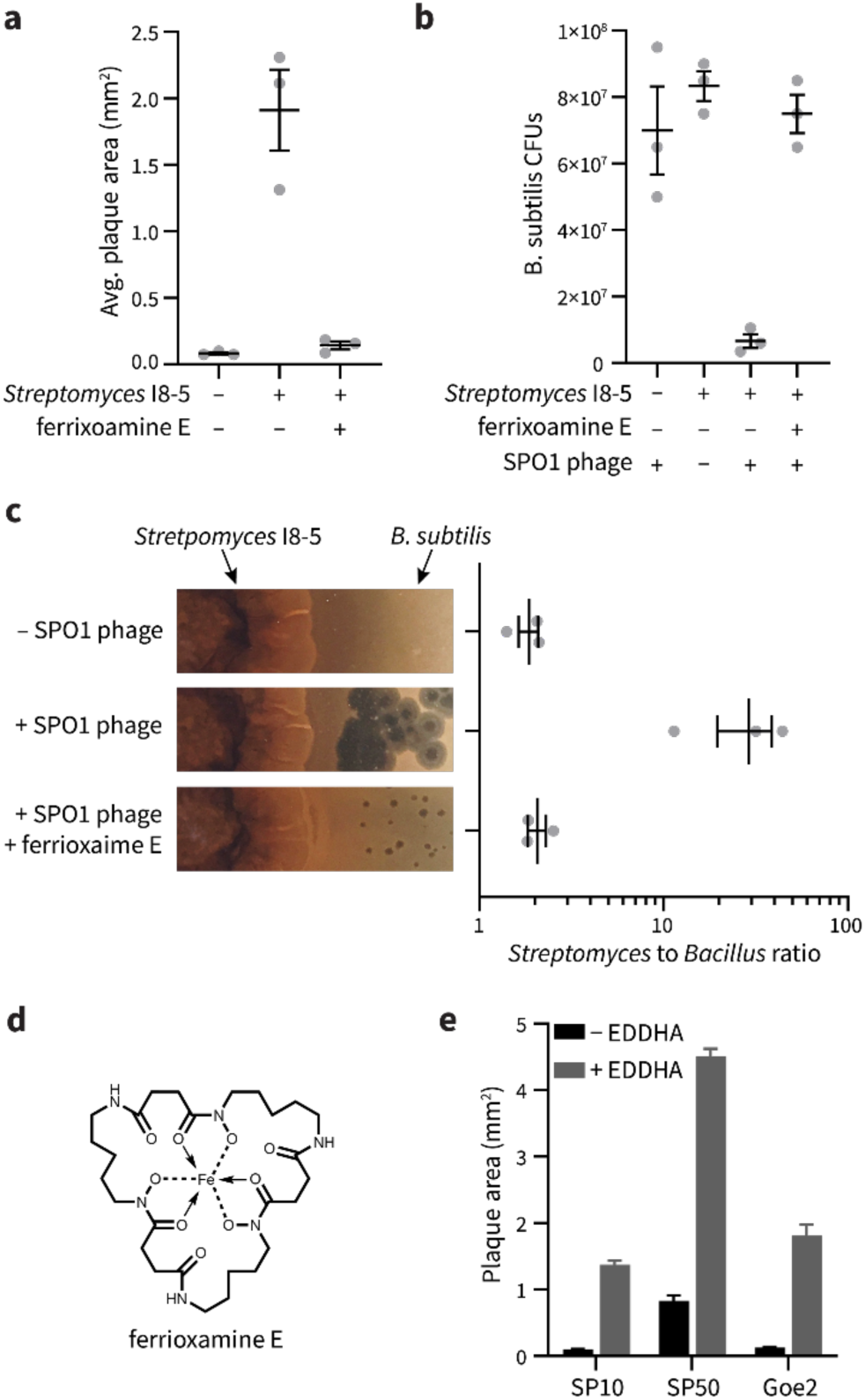
Coelichelin helps *Streptomyces* to outcompete *B. subtilis*. **(a)** The average plaque areas of SPO1 on *B. subtilis* alone, and *B. subtilis* neighboring *Streptomyces* I8-5 colony with or without ferrioxamine E as excess iron source. At least eight plaques were selected for each replicate. Data are represented as the average ± SEM from three independent biological replicates. Circles show the values of each biological replicate. **(b)** The colony forming units of *B. subtilis* measured in the presence of phages, neighboring *Streptomyces* I8-5 colony with and without phages, and neighboring *Streptomyces* I8-5 colony with ferrioxamine E as excess iron source in the presence of phages. Data are represented as the average ± SEM from three independent biological replicates. Circles show the values of each biological replicate. **(c)** *Streptomyces* to *B. subtilis* ratio calculated from colony forming units measured after 2 days of co-culture. Data are represented as the average ± SEM from three independent biological replicates. Circles show the values of each biological replicate. **(d)** The chemical structure of ferrioxamine E. **(e)** The plaquing promotion effect of EDDHA (6 mM) for phages SP10, SP50, and Goe2 on *B. subtilis*. Water was used as the −EDDHA control. Data are represented as the average ratio ± SEM calculated from at least four individual plaques of each condition.

Finally, we asked if the facilitated phage predation was specific to only the SPO1 phage, or if it could be generalized to a wide range of phages that would encounter mixed *B. subtilis* + *Streptomyces* communities in nature. We tested three other *Bacillu*s phages (SP10, SP50, and Goe2),^38, 39^ and the virulence of all three was substantially increased by iron sequestration (Fig. 5e). Therefore, iron sequestration by competing microbes may broadly sensitize *B. subtilis* to a variety of phages in nature.

## Discussion

We discovered that a bacterial strain gains a competitive advantage over neighboring bacteria by producing a secondary metabolite that sensitizes its competitor to phage predation. In our culture conditions, the metabolite/phage synergy switched the outcome of the bacterial competition to strongly favor the metabolite producer. The secondary metabolite is the known *Streptomyces*-produced siderophore coelichelin, which sequesters iron away from *B. subtilis* and promotes phage infection by inhibiting sporulation. These findings reveal a new mechanism by which siderophores can shape microbial competition through bacteria-phage ecology.

Siderophores are primarily believed to provide a means for the producing organism to acquire scarce iron ions to supply their own metalloenzymes.^6, 20^ Beyond iron acquisition, siderophores can benefit their producers in other ways that justify the maintenance of their biosynthetic genes. Notably, siderophores can inhibit the growth of neighbors, allowing the producing microbe to displace its competitors.^6, 40^ Our results also show that siderophores can block sporulation of competing bacteria, which could be beneficial in fluctuating environments. For example, by excluding spores of a competitor, the siderophore-producer’s spores would revive without competition when conditions become optimal for growth and reproduction.^41–43^ Finally, our work reveals a potential fourth benefit of siderophore production: microbes can sensitize competing bacteria to lytic phages. Since iron sequestration sensitizes *B. subtilis* to several natural phages, this phage-promoting effect may benefit the multitude of siderophore-producing bacteria and fungi^20^ that compete with *B. subtilis* and its endospore-forming relatives.

Beyond microbe-microbe competition, multicellular hosts also leverage iron sequestration to stall the growth of pathogens by producing molecules like transferrin and lactoferrin.^44^ Therefore, host-induced iron sequestration may also sensitize certain endospore-forming bacteria to lytic phages (either ones endogenously present or those administered therapeutically).

Although our studies focused on soil *Streptomyces* and the model soil bacterium *B. subtilis*, endosporulation is a characteristic trait of many bacteria in the Bacillota (Firmicutes) phylum.^45^ These endospore-forming bacteria are not only abundant in soil and aquatic sediments, but they are also important members of host-associated microbiomes—including some common intestinal pathogens and mutualistic taxa in humans.^46^ Therefore, it is plausible that secondary metabolites sensitize diverse Bacillota (Firmicutes) in varied environments to phage infection via sporulation inhibition. In fact, secondary metabolites other than siderophores have also been shown to inhibit sporulation. For example, a common signal molecule used for quorum sensing, autoinducer-2, inhibits sporulation in *Bacillus velezensis*.^34^ Furthermore, the bacterial macrocycle, fidaxomicin, inhibits *Clostridioides difficile* sporulation.^47^ Therefore, microbial siderophores, other microbial secondary metabolites, and even host-produced molecules may sensitize competing bacteria to phage infection in natural communities.

In conclusion, we found an example that a natural product, coelichelin, gives its producer an advantage by sensitizing its competitors to phages. Despite a rich history of studies on the “chemical warfare” waged between microbes via natural products, little emphasis has been placed on how phage predation intersects with microbial secondary metabolites. This work reveals that microbial natural products do not just directly inhibit the fitness of microbial competitors, but these molecules can also sensitize competitors to lytic phages. Although the extent of this phenomenon in nature is yet to be seen, it may shape the microbial ecology of both environmental and host-associated ecosystems. Also, much like society has leveraged microbial competition to discover life-saving antimicrobials, phage-promoting natural products may also prove useful one day as co-administered adjuvants in phage-based interventions.

## Methods

### Strains and growth conditions

The strains and bacteriophages used in this study are listed in Table S1. All chemicals used in this study are listed in Table S2. All primers used in this study are listed in Table S3. *B. subtilis* strains were routinely grown in LB broth at 37 ℃ and 220 rpm. When appropriate, antibiotics were used at the following concentrations: 7 μg/mL kanamycin and 1 μg/mL erythromycin. *Streptomyces* strains were routinely grown in ISP2 media (4 g/L yeast extract, 10 g/L malt extract, and 4 g/L dextrose) at 30 ℃ and 220 rpm.

### Bacteriophage lysate preparation

To prepare the host culture, an overnight culture of *B. subtilis* RM125 WT was sub-cultured 1:100 into 4 mL LB + 0.1 mM MnCl_2_ + 5 mM MgCl_2_ + 5 mM CaCl_2_. The culture was incubated at 37 ℃ and 220 rpm for 4 hours until the OD_600nm_ reached 0.2. About 1×10^3^ plaque forming units (PFUs) of bacillus phage were added to the culture. The phage infected culture was incubated at 37 ℃ and 220 rpm until bacterial cells were lysed and the culture turned clear. The phage lysate was filtered through a 0.2 µm polyethersulfone filter and stored at 4 ℃.

### Binary-interaction screening

To prepare plates with library bacteria, 5 μL of the frozen spore stock of each bacteria strain in our library was suspended in 50 μL of ISP2 medium. Then 8 μL of the spore suspension was spotted at the center of a ISP2 + 1.5% agar plate. The inoculated plates were incubated at 30 ℃ for 10 days to allow the library bacteria to grow and secrete their metabolites into the plate. To test the influence of the metabolites on phage infectivity, an overnight culture of *B. subtilis* RM125 WT was diluted 1:10 into 5 mL fresh LB broth and then ∼1,000 PFUs of SPO1 phages were added into the medium. The mixture of bacteria and phages was poured around the central colony formed by the library bacteria. Bacteria and phages were allowed 10 mins to soak the plate, and then the bacteria dilution was removed by pipet. The plate was dried under room temperature in biosafety cabinet. The plate was incubated at 37 ℃ overnight and the plaques formed by SPO1 was examined the next day.

### 16S rRNA sequencing of I8-5

A single colony of I8-5 was inoculated into 4 mL fresh ISP2 medium and incubated at 30 ℃, 220 rpm for 4 days. The genomic DNA was extracted from 1 mL liquid culture using Promega Wizard Genomic DNA Purification Kit (#1120). The 16S rRNA region of I8-5 genome was amplified by PCR using 16S_F and 16S_R primers. Sanger sequencing result of its 16S rRNA with 16S_F primer was available on NCBI (accession number: OR902106). The 16S rRNA sequence of I8-5 aligns well with species in *Streptomyces* genus. (Supplementary file 1)

### Coelichelin biosynthetic gene cluster identification in I8-5

The library of the extracted genomic DNA was prepared by Illumina Nextera XT DNA Library Prep Kit protocol (# FC-131-1096) and analyzed by Agilent D1000 ScreenTape. The libraries were pooled and loaded on a NextSeq 1000/2000 P2 Reagents (100-cycles) v3 flow cell (#20046811) configured to generate paired end reads. The demultiplexing of the reads was performed using bcl2fastq, version 2.20.0. Reads were adapter trimmed and quality filtered using Trimmomatic^48^ 0.38 with the cutoff threshold for average base quality score set at 20 over a window of 3 bases requiring a minimum trimmed read length of 20 bases (parameters: LEADING:20 TRAILING:20 SLIDINGWINDOW:3:20 MINLEN:20). The cleaned reads were assembled using SPAdes^49^ version 3.15.4 with default parameters. The assembly was annotated using prokka^50^ version 1.12 employing a sequence training set prepared from protein sequences obtained from 431 publicly available Streptomyces assemblies (parameters: --minpid 70 --usegenus --hmmlist TIGRFAM,CLUSTERS,Pfam,HAMAP). The coelichelin biosynthetic gene cluster was identified by antiSMASH 7.0.^51^ The genomic sequences are available on NCBI (accession number: JAYMFC000000000).

### Collection of I8-5 supernatant

To revive the spores of I8-5, 5 μL of the I8-5 frozen spore stock was streaked out on a ISP2 + 1.5% agar plate. The plate was incubated at 30℃ for 3 days until colonies formed. A colony was inoculated into 4 mL fresh ISP2 medium and incubated at 30 ℃, 220 rpm for 4 days. After the incubation, 1 mL of the culture was added into 100 mL of ISP2 medium and grown for another 11 days to allow metabolite production. To harvest the metabolites in the supernatant, the bacteria cells in the culture were pelleted at 4,820×g for 20 min and discarded. The supernatant was lyophilized and stored at -20 ℃ until ready to use.

### Phage promotion activity test

An overnight culture of *B. subtilis* RM125 WT was diluted 1:10 into 5 mL fresh ISP2 + 0.1 mM MnCl_2_ + 5 mM MgCl_2_, and poured onto an ISP2 + 0.1 mM MnCl_2_ + 5 mM MgCl_2_ + 1.5% agar plate. Bacteria were allowed 10 min to attach to the plate, and then the unattached bacteria were removed. The plate was dried under room temperature in biosafety cabinet. To test the phage promotion effect, 2 μL of compound was spotted on top of the bacterial lawn. After the compound dried, incubated the plate at 37 ℃ for 1 h. Then 5 μL of SPO1 phage lysate (∼10 PFUs) was spotted on top of the compound treated area. After the phage lysate dried, the plate was incubated at 37 ℃ overnight and the plaques formed by SPO1 was examined the next day.

### Fractionation of I8-5 supernatant using reversed-phase chromatography

The lyophilized supernatant was dissolved into a small amount of water as a concentrated sample. The concentrated supernatant was further separated on a Phenomenex Synergi 4 μm Hydro-RP 80 Å column (250 × 10 mm) using an Agilent 1260 Infinity II HPLC system. The mobile phase A was water + 0.01 % (v/v) formic acid and the mobile phase B was acetonitrile + 0.01 % (v/v) formic acid. The flow rate was kept at 3 mL·min^-1^ and the gradient was as follows: 0% B (0–5 min), increase to 20% B (5–6 min), 20% B for (6–11 min), increase to 80% B (11–31 min), increase to 100% B (31–32 min), 100% B (32–37 min), decrease to 0% B (37–38 min), 0% B (38–43 min). Eluted fractions were collected every 30 seconds and dried in vacuo. Each dried fraction was redissolved into 2 μL DMSO and spotted on a lawn of *B. subtilis* RM125 WT infected with ∼1000 PFUs of SPO1 phages to test the phage promotion activity. The fraction eluting at 12.0∼12.5 min is active and labeled as “active fraction 1”. The composition of “active fraction 1” was analyzed on a Phenomenex Synergi 4 μm Hydro-RP 80 Å column (250×4.6 mm) using an Agilent 1260 Infinity II HPLC system coupled to a mass spectrometer Agilent InfinityLab LC/MSD XT. The analysis was performed at a flow rate of 0.7 mL·min^–1^. The mobile phase and separation gradient were the same as described above.

### Fractionation of I8-5 supernatant using cation-exchange and reversed-phase chromatography

The lyophilized supernatant was dissolved into 10 mL water and added 2% (v/v) formic acid to adjust the pH to 2.02. The supernatant was loaded on a Waters Oasis MCX column (#186000255). The column was then eluted with water + 2% (v/v) formic acid, methanol, and methanol + 5% (v/v) ammonium hydroxide. The eluates were dried in vacuo and redissolved into water as a 100 mg/mL solution. 2 μL of each redissolved fraction was spotted on a lawn of *B. subtilis* RM125 WT infected with ∼1000 PFUs of SPO1 phages to test the phage promotion activity. The methanol + 5% (v/v) ammonium hydroxide eluate was active and subjected to separation on a Phenomenex Synergi 4 μm Hydro-RP 80 Å column (250 × 10 mm) using an Agilent 1260 Infinity II HPLC system. The mobile phase A was water + 0.1 % (v/v) formic acid and the mobile phase B was acetonitrile + 0.1 % (v/v) formic acid. The flow rate was kept at 3 mL·min^–1^ and the gradient was as follows: 10% B (0–10 min), increase to 50% B (10–30 min), increase to 100% B for (30–31 min), 100% B (31–38 min), decrease to 10% B (38–39 min), 10% B (39–44 min). Eluted fractions were collected every 30 seconds and dried in vacuo. Each dried fraction was redissolved into 2 μL DMSO tested for phage promotion effect as described above. The fraction eluting at 3.5∼4.0 min is active and labeled as “active fraction 2”. The composition of “active fraction 2” was analyzed on a Phenomenex Synergi 4 μm Hydro-RP 80 Å column (250×4.6 mm) using an Agilent 1260 Infinity II HPLC system coupled to a mass spectrometer Agilent InfinityLab LC/MSD XT. The analysis was performed at a flow rate of 0.7 mL·min^–1^. The mobile phase and separation gradient were the same as described above.

### Tandem mass spectrometry (MS/MS) analysis of 566.2867 [M+H]^+^

High-resolution electrospray ionization (HR-ESI) mass spectra with collision-induced dissociation (CID) MS/MS were obtained using a Waters Synapt G2S QTOF. Data-dependent acquisition was employed to fragment the top three masses in each scan. The “active fraction 2” was separated on a Phenomenex Synergi 4 μm Hydro-RP 80 Å column (250×4.6 mm). The mobile phase A was water + 0.1 % (v/v) formic acid and the mobile phase B was acetonitrile + 0.1 % (v/v) formic acid. The flow rate was kept at 0.7 mL·min^–1^and the gradient was as follows: 0% B (0–20 min), increase to 20% B (20–21 min), increase to 40% B for (21–31 min), increase to 100% B (31–32 min), 100% B (32–42 min), decrease to 0% B (42–43 min), 0% B (43–48 min). 566.2867 [M+H]^+^ eluted at 17.2∼18.2 min.

### Isolation of coelichelin from I8-5 supernatant

The protocol was adapted from Challis et al.^17^ The lyophilized I8-5 supernatant (1.8533 g) was dissolved in 5 mL of water. FeCl_3_ was added to the supernatant (final concentration 40 mM) to generate Fe-coelichelin complex. The reaction mixture was centrifuged at 16,000×g for 5 min and the precipitates were discarded. The supernatant was separated on a Phenomenex Luna 10 μm Hydro-RP 100 Å column (250×21.2 mm) using an Agilent 1260 Infinity II HPLC system. The mobile phase A was water + 10 mM NH_4_HCO_3_ (pH=8.01) and the mobile phase B was methanol. The flow rate was kept at 10 mL·min^–1^ and the gradient was as follows: 5% B (0–20 min), increase to 90% B (20–21 min), 20% B for (21–31 min), decrease to 5% B (31–32 min), 5% B (32–42 min). Fe-coelichelin eluted at 9.1∼10.4 min and was collected by monitoring the absorbance at 435 nm. The collected Fe-coelichelin was concentrated in vacuo, lyophilized, and obtained as an orange solid.

The obtained Fe-coelichelin (45.9 mg) was dissolved into 74 mL water. The ferric iron was removed from Fe-coelichelin complex by mixing the Fe-coelichelin solution with 74 mL 100 mM 8-hydroxyquinoline in methanol. The reaction was stirred for 30 min at room temperature. The Fe-8-hydroxyquinoline complex was removed by extracting the aqueous phase using 50 mL dichloromethane 3 times. The aqueous phase was concentrated in vacuo and separated on a Phenomenex Synergi 4 μm Hydro-RP 80 Å column (250×10 mm) using an Agilent 1260 Infinity II HPLC system. The mobile phase A was water + 0.1 % (v/v) formic acid and the mobile phase B was acetonitrile + 0.1 % (v/v) formic acid. The flow rate was kept at 3 mL·min^–1^ and the gradient was as follows: 0% B (0–10 min), increase to 5% B (10–11 min), 5% B for (11–21 min), increase to 100% B (21–22 min), 100% B (22–32 min), decrease to 0% B (32–33 min), 0% B (33–43 min). Apo-coelichelin eluted at 17.7∼17.9 min and was collected by monitoring the absorbance at 210 nm. The collected apo-coelichelin was concentrated in vacuo, lyophilized, and obtained as a white solid. The high-resolution electrospray ionization mass spectrometry (HR-ESI-MS) data of apo-coelichelin was obtained on a Thermo Scientific Finnigan LTQ Orbitrap XL Mass Spectrometer equipped with a nano-electrospray ionization source operated in positive ionization mode. HR-ESI-MS (positive-ion mode): *m/z* 566.2776 [M+H]^+^ (calcd for C_21_H_40_N_7_O_11_^+^: 566.2780)

### Coelichelin purity check by LC-MS

The purified coelichelin was analyzed on a Phenomenex Synergi 4 μm Hydro-RP 80 Å column (250×4.6 mm) using an Agilent 1260 Infinity II HPLC system coupled to a mass spectrometer Agilent InfinityLab LC/MSD XT. The mobile phase A was water + 0.1 % (v/v) formic acid and the mobile phase B was acetonitrile + 0.1 % (v/v) formic acid. The flow rate was kept at 0.7 mL·min^–1^and the gradient was as follows: 0% B (0–10 min), increase to 5% B (10–30 min), increase to 100% B for (30–31 min), 100% B (31–41 min), decrease to 0% B (41–42 min), 0% B (42–52 min).

### Preparation of Ga-coelichelin

Coelichelin (10 mg) was dissolved in 400 μL of water and reacted with 26.5 mg of Ga_2_(SO_4_)_3_ in 400 μL of water. The reaction was performed at room temperature for 30 mins and subjected to separation on a Phenomenex Synergi 4 μm Hydro-RP 80 Å column (250×10 mm) using an Agilent 1260 Infinity II HPLC system. The mobile phase A was water + 0.1 % (v/v) formic acid and the mobile phase B was acetonitrile + 0.1 % (v/v) formic acid. The flow rate was kept at 3 mL·min^–1^and the gradient was as follows: 0% B (0–10 min), increase to 100% B (10–11 min), 100% B for (11–21 min, decrease to 0% B (21–22 min), 0% B (22–32 min). Ga-coelichelin eluted at 4.6∼5.0 min and was collected by monitoring the absorbance at 210 nm. The collected Ga-coelichelin was concentrated in vacuo, lyophilized, and obtained as a white solid. HR-ESI-MS (positive-ion mode): *m/z* 632.1794 [M+H]^+^ (calcd for C_21_H_37_GaN_7_O_11_^+^: 632.1801). ^1^H and TOCSY (mixing time of 60 ms) NMR spectra were obtained on a Varian 600 MHz Inova NMR spectrometer. ^13^C, DQF-COSY, HSQC, and HMBC NMR spectra were obtained on a Bruker 500 MHz Avance Neo NMR spectrometer.

### Iron complementation experiment

An overnight culture of *B. subtilis* RM125 WT was diluted 1:10 into 5 mL fresh ISP2 + 0.1 mM MnCl_2_ + 5 mM MgCl_2_, and poured onto an ISP2 + 0.1 mM MnCl_2_ + 5 mM MgCl_2_ + 1.5% agar plate. Bacteria were allowed 10 mins to soak the plate, and then the bacteria dilution was removed by pipet. The plate was dried under room temperature in a biosafety cabinet. 2 μL of compound was spotted as a small circle on top of the bacterial lawn. After the compound dried, 2 μL of FeSO_4_ aqueous solution was spotted on top of the compound-treated area. After the FeSO_4_ solution dried, the plate was incubated at 37 ℃ for 1 hour. Then 5 μL of SPO1 phage lysate (∼10 PFUs) was spotted on top of the compound treated area. After the phage lysate dried, the plate was incubated at 37℃ overnight and the plaques formed by SPO1 were examined the next day.

### Quantification of phage reproduction from individual plaques

An overnight culture of *B. subtilis* RM125 WT was diluted 1:10 into 5 mL fresh ISP2 + 0.1 mM MnCl_2_ + 5 mM MgCl_2_, and poured onto an ISP2 + 0.1 mM MnCl_2_ + 5 mM MgCl_2_ + 1.5% agar plate. Bacteria were allowed 10 mins to soak the plate, and then the bacteria dilution was removed by pipet. The plate was dried under room temperature in biosafety cabinet. 2 μL of 6 mM EDDHA or water was spotted as a small circle on top of the bacterial lawn. After the compound dried, the plate was incubated at 37 ℃ for 1 hour. Then 5 μL of SPO1 phage lysate (10∼40 PFUs) was spotted on top of the compound treated area. After the phage lysate dried, the plate was incubated at 37℃ for 2 days. The number of plaques formed in each phage spot were enumerated and the average plaque areas were measured for each phage spot. All the plaques in one phage spot were pooled by carving out the agar with the plaques and resuspended in 5 ml phage buffer (10 mM Tris, 10 mM MgSO_4_, 4g/L NaCl, pH=7.5). The suspension was vortexed at the highest speed for 20 seconds to allow phages to fully detach from the agar. The PFUs of the pooled plaques were quantified by the small drop plaque assay.^52^ For each individual phage spot, the average PFUs per plaque was calculated using the following equation:

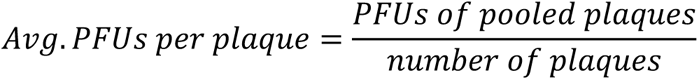

### Sporulation quantification

An overnight culture of *B. subtilis* RM125 WT was diluted 1:10 into 5 mL fresh ISP2 + 0.1 mM MnCl_2_ + 5 mM MgCl_2_, and poured onto an ISP2 + 0.1 mM MnCl_2_ + 5 mM MgCl_2_ + 1.5% agar plate. Bacteria were allowed 10 mins to soak the plate, and then the bacteria dilution was removed by pipet. The plate was dried under room temperature in biosafety cabinet. 2 μL of compound or water (control) was spotted as a small circle on top of the bacterial lawn. After the compound dried, the plate was at 37 ℃ for 16 hours. Then ∼1cm^2^ area of bacteria was scraped off the plate and resuspended in 200 μL water. The cell suspension was heated at 85 ℃ for 15 mins to kill non-sporulated cells. Then the spores in the heat-treated cell suspension were quantified by measuring the colony forming units.

### Generation of sporulation mutants

The gene knockout mutants in *B. subtilis* 168 were purchased from Bacillus Genomic Stock Center. The mutation was then transferred to *B. subtilis* RM125 using SPP1-mediated generalized phage transduction.^53, 54^ The SPP1 phage lysate was obtained from the *B. subtilis* 168 knockout donor strain as described above and stored at 4 ℃ until ready to use. A single colony of the recipient *B. subtilis* RM125 was inoculated into 10 mL LB + 10 mM CaCl_2_. The recipient culture was incubated at 37 ℃ and 220 rpm for 4 hours. For phage transduction, 950 μL of the recipient culture was mixed with 50 μL of donor SPP1 lysate, and incubated at 37 ℃ for 10 mins to allow phage adsorption. Then the infected culture was transferred into 9 mL of prewarmed LB + 20 mM sodium citrate and incubated at 37 ℃ for another 10 mins. The cells were pelleted at 4000×g for 5 min and plated onto an LB + 20 mM sodium citrate + 1.5% agar plate with appropriate antibiotics. The plates were incubated at 37 ℃ overnight and the mutant colonies were re-streaked twice on LB + 20 mM sodium citrate + 1.5% agar plate with appropriate antibiotics to clean out phages. The knockout mutation was validated by PCR with primers reported by Gross et al.^55^ The sporulation mutants were verified to not produce spores by the sporulation quantification experiment described above.

### Streptomyces sp. I8-5 and B. subtilis competition

To inoculate plates with *Streptomyces*, 5 μL of the frozen spore stock of I8-5 was suspended in 50 μL of ISP2 + 0.1 mM MnCl_2_ + 5 mM MgCl_2_. Then 8 μL of the spore suspension was spotted at the center of an ISP2 + 0.1 mM MnCl_2_ + 5 mM MgCl_2_ + 1.5% agar plate. The plates were incubated at 30℃ for 22 days to allow I8-5 colony to grow and secrete coelichelin into the plate. To inoculate the *B. subtilis* next to *Streptomyces*, an overnight culture of *B. subtilis* RM125 WT was diluted 1:10 into 5 mL fresh ISP2 + 0.1 mM MnCl_2_ + 5 mM MgCl_2_, and poured around the I8-5 colony. *B. subtilis* cells were allowed 10 mins to soak the plate, and then the bacteria dilution was removed by pipet. The plate was dried under room temperature in biosafety cabinet. 2 μL of water or ferrioxamine E was spotted as a small circle on top of the *B. subtilis* lawn. After the spotted solution dried, the plate was incubated at 37 ℃ for 1 hour. Then 5 μL of SPO1 phage lysate (∼10 PFUs) was spotted on top of the compound treated area. After the phage lysate dried, the plate was incubated at 37℃ for two days. The average plaque area was measured for each phage spot. The *B. subtilis* lawn and *Streptomyces* colony were carved out and resuspended in 5 ml LB broth and 5 ml ISP2 medium respectively. The cell suspensions were vortexed at the highest speed for 20 seconds to allow bacteria cells to fully detach from the agar. The colony forming units of *B. subtilis* cell suspensions were quantified by plating serial dilutions of the cell suspension on LB + 1.5% agar plates. The colony forming units of *Streptomyces* cell suspensions were quantified by plating serial dilutions of the cell suspension on ISP2 + 10 ug/ml nalidixic acid + 1.5% agar plates.

## Acknowledgments

We thank Andreea Măgălie (Georgia Institute of Technology) and Joshua Weitz (University of Maryland) for helpful discussions. We thank the Bacillus Genomic Stock Center (Ohio State University), the Félix d’Hérelle Reference Center for Bacterial Viruses (University of Laval), and Robert Hertel (University of Goettingen) for providing bacteria and phages. We thank E. M. Nolan (Massachusetts Institute of Technology) for providing enterobactin. The research was supported by a research starter grant from the American Society of Pharmacognosy to J.P.G. and a National Science Foundation CAREER award (IOS-2143636) to J.P.G. Research support was also provided by the National Science Foundation (DEB-1934554 to J.T.L., D.A.S.; DBI-2022049 to J.T.L.), the US Army Research Office (W911NF-22-1-0014, W911NF-22-S-0008 to J.T.L.), and the National Aeronautics and Space Administration (80NSSC20K0618 to J.T.L.). Z.Z. was supported in part by the John R. and Wendy L. Kindig Fellowship. K.J.P. and the Laboratory for Biological Mass Spectrometry were supported by the Indiana University Precision Health Initiative. The 500 MHz NMR and 600 MHz spectrometer of the Indiana University NMR facility were supported by NSF grant CHE-1920026, and the Prodigy probe was purchased in part with support from the Indiana Clinical and Translational Sciences Institute funded, in part, by NIH Award TL1TR002531.

## Author Contributions

Conceptualization, Z.Z., J.P.G.; Methodology, Z.Z., D.S., J.P.G.; Investigation, Z.Z., C.Z., K.J.P., R.P.; Writing – Original Draft, Z.Z., J.P.G.; Writing – Review & Editing, Z.Z., C.Z., D.S., J.T.L., J.P.G.; Visualization, Z.Z., J.P.G.; Supervision, J.T.L., J.P.G.; Funding Acquisition, J.T.L., J.P.G.

---

**Extended data Fig. 1.**
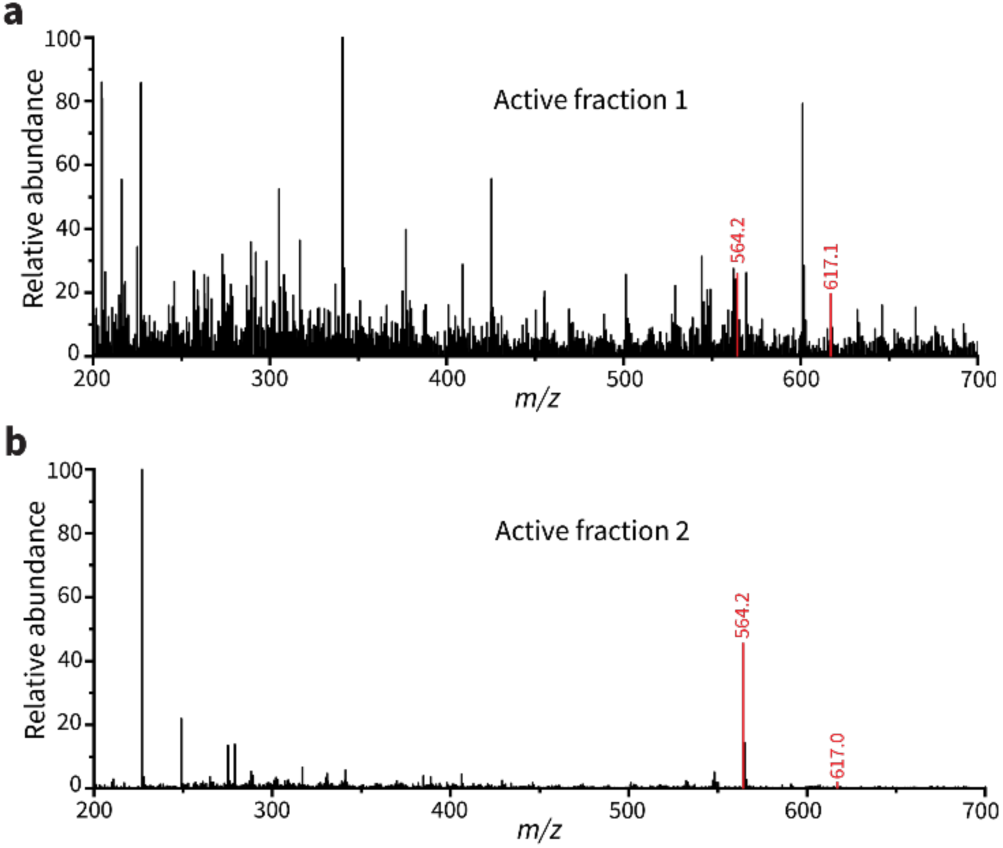
The negative mode electrospray ionization MS spectra of active fraction 1 (a) and active fraction 2 (b). The shared peaks are highlighted red.

**Extended Data Fig. 2.**
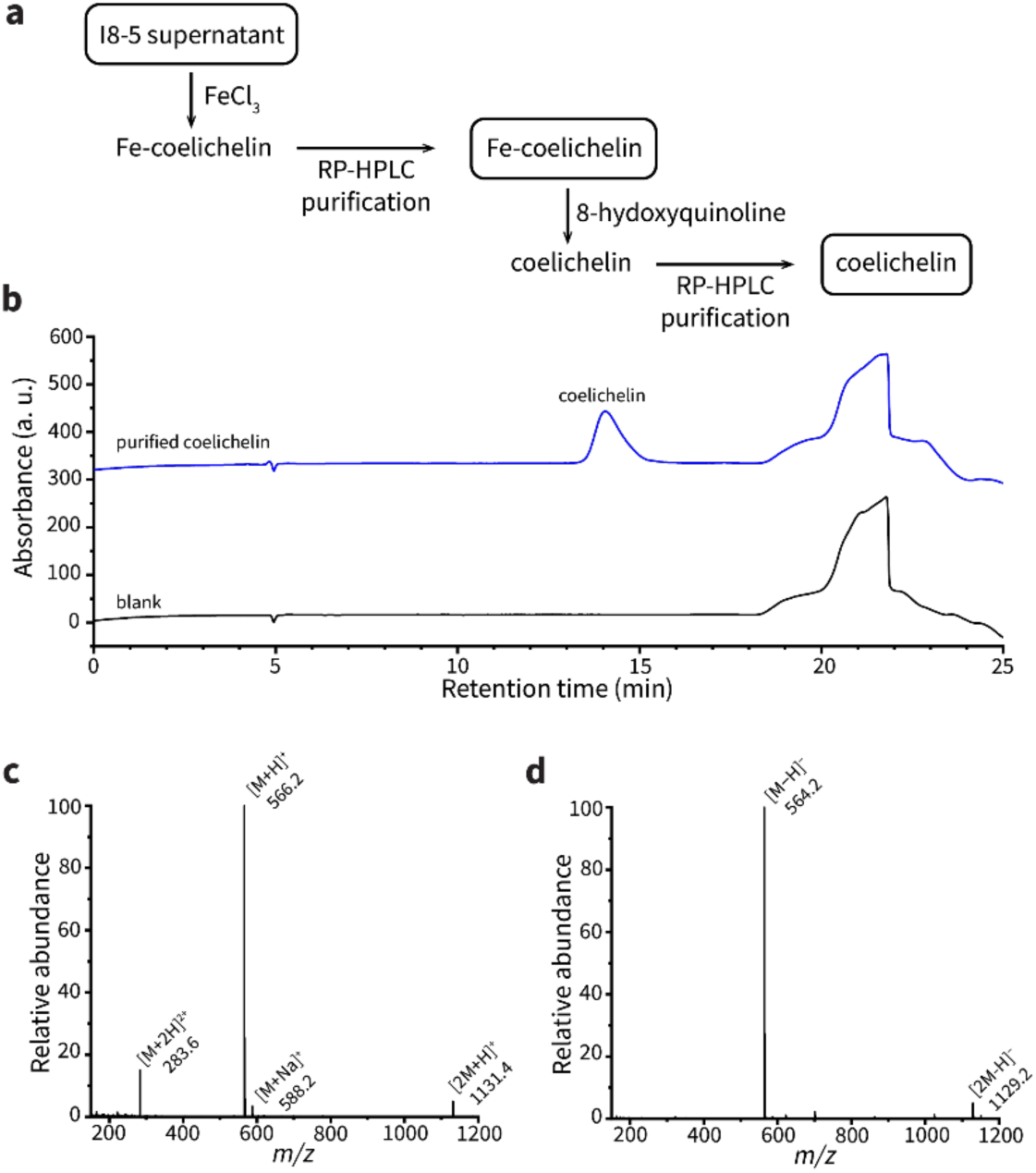
Coelichelin isolation from I8-5 supernatant. (**a**) Isolation scheme. (**b**) UV chromatogram at 210 nm. Water was used as the blank. (c) The averaged MS spectrum at positive mode between retention time 13.5∼14.8 min. (d) The averaged MS spectrum at negative mode between retention time 13.5∼14.8 min. M represents coelichelin.

**Extended Data Fig. 3.**
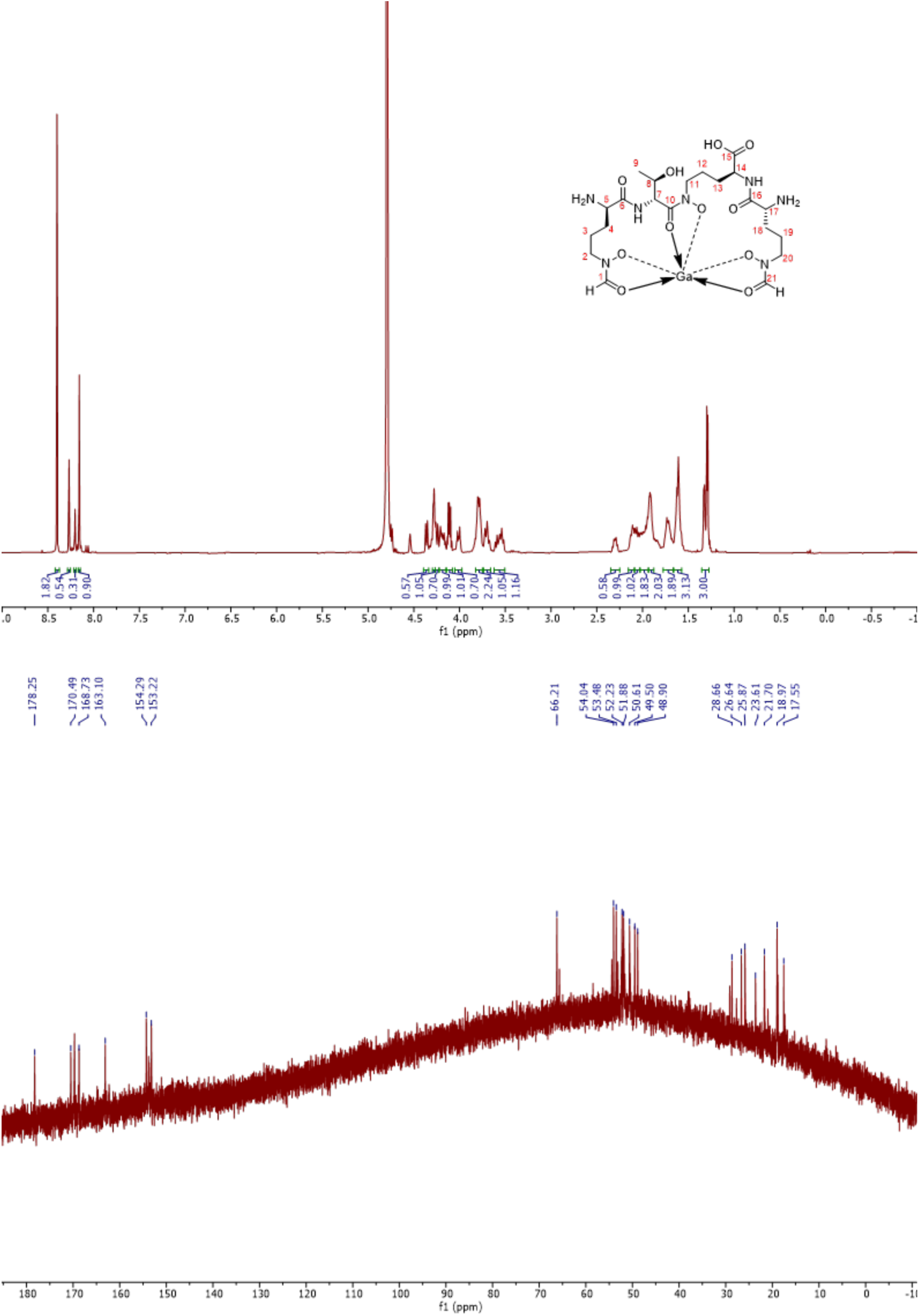
^1^H NMR spectrum (600 MHz, top) and ^13^C NMR spectrum (125 MHz, bottom) of Ga-coelichelin in D_2_O.

**Extended Data Table 1.** Summary of 1H (600 MHz) and ^13^C (125 MHz) NMR spectral data of Ga-coelichelin in D_2_O.

| Position | δ<sub>C</sub> | δ<sub>H</sub> (mult., J in Hz) | Position | δ<sub>C</sub> | δ<sub>H</sub> (mult., J in Hz) |
| --- | --- | --- | --- | --- | --- |
| 1 | 153.22 | 8.16 (1H, s) | 12 | 21.70 | 1.91 (1H, m) |
| 2 | 50.61 | 3.78 (1H, m) |  |  | 1.62 (1H, m) |
|  |  | 3.56 (1H, m) | 13 | 23.61 | 1.98 (2H, m) |
| 3 | 17.55 | 1.72 (1H, m) | 14 | 54.04 | 4.25 (1H, d, 12.2) |
|  |  | 1.62 (1H, m) | 15 | 178.25 | - |
| 4 | 25.87 | 2.12 (1H, m) | 16 | 170.49 | - |
|  |  | 1.62 (1H, m) | 17 | 52.23 | 4.36 (1H, d, 11.0) |
| 5 | 51.88 | 4.28 (1H, m) | 18 | 28.66 | 2.30 (1H, dt, 16.2, 9.0) |
| 6 | 168.73 | - |  |  | 1.91 (1H, m) |
| 7 | 53.48 | 4.79<sup>a</sup> (1H) | 19 | 26.64 | 2.06 (1H, m) |
| 8 | 66.21 | 4.11 (1H, dp, 9.2, 6.3) |  |  | 1.72 (1H, m) |
| 9 | 18.97 | 1.29 (3H, d, 6.3) | 20 | 48.90 | 4.01 (1H, dt, 14.8, 6.0) |
| 10 | 163.10 | - |  |  | 3.78 (1H, m) |
| 11 | 49.50 | 4.19 (1H, m) | 21 | 154.29 | 8.27 (1H, s) |
|  |  | 3.70 (1H, m) |  |  |  |

<sup>a</sup>Proton signal was masked by solvent residual

