## Supplemental Tables S1-S3 and Figures S1-S4 for "Bacterium secretes chemical inhibitor that sensitizes competitor to bacteriophage infection"

### Table of Contents

Supplementary Table S1. Strains and bacteriophages used in this study..................................... S3
Supplementary Table S2. Chemical used in this study ................................................................ S5
Supplementary Table S3. Primers used in this study .................................................................... S6
Supplementary Figure S1. DQF-COSY NMR spectrum of Ga-coelichelin in $D_2O$ (500 MHz). . S7
Supplementary Figure S2. TOCSY (mixing time of 60 ms) NMR spectrum of Ga-coelichelin in $D_2O$ (600 MHz)........................................................................................................................ S8
Supplementary Figure S3. HSQC NMR spectrum of Ga-coelichelin in $D_2O$. ............................ S9
Supplementary Figure S4. HMBC NMR spectrum of Ga-coelichelin in $D_2O$. .......................... S10
Reference ........................................................................................................................ S11

**Supplementary Table S1. Strains and bacteriophages used in this study**

| Bacterial strains | Description | Source |
|---|---|---|
| ***B. subtilis* RM125 WT** | Wild type strain | Bacillus Genetic Stock Center #1A253 |
| ***B. subtilis* 168 Δ*spo0A*::Kan$^R$** | Δ*spo0A* donor, kanamycin-resistant | Bacillus Genetic Stock Center #BKK24220 |
| ***B. subtilis* 168 Δ*spo0B*::Kan$^R$** | Δ*spo0B* donor, kanamycin-resistant | Bacillus Genetic Stock Center #BKK27930 |
| ***B. subtilis* 168 Δ*spo0F*::Kan$^R$** | Δ*spo0F* donor, kanamycin-resistant | Bacillus Genetic Stock Center #BKK37130 |
| ***B. subtilis* 168 Δ*spoIIE*::Erm$^R$** | Δ*spoIIE* donor, erythromycin-resistant | Bacillus Genetic Stock Center #BKE00640 |
| ***B. subtilis* 168 Δ*spoIIA*::Kan$^R$** | Δ*spoIIA* donor, kanamycin-resistant | Bacillus Genetic Stock Center #BKK23470 |
| ***B. subtilis* 168 Δ*sigF*::Erm$^R$** | Δ*sigF* donor, kanamycin-resistant | Bacillus Genetic Stock Center #BK23450 |
| ***B. subtilis* RM125 Δ*spo0A*::Kan$^R$** | *spo0A* knockout mutant, kanamycin-resistant | This study |
| ***B. subtilis* RM125 Δ*spo0B*::Kan$^R$** | *spo0B* knockout mutant, kanamycin-resistant | This study |
| ***B. subtilis* RM125 Δ*spo0F*::Kan$^R$** | *spo0F* knockout mutant, kanamycin-resistant | This study |
| ***B. subtilis* RM125 Δ*spoIIE*::Erm$^R$** | *spoIIE* knockout mutant, erythromycin-resistant | This study |
| ***B. subtilis* RM125 Δ*spoIIA*::Kan$^R$** | *spoIIA* knockout mutant, kanamycin-resistant | This study |
| ***B. subtilis* RM125 Δ*sigF*::Erm$^R$** | *sigF* knockout mutant, kanamycin-resistant | This study |
| ***Streptomyces* I8-5** | Isolated from standard brown-capped gilled mushroom, Bloomington, IN | Lab collection |

**Bacteriophages**

| | | |
|---|---|---|
| **SPO1** | Bacillus phage | Bacillus Genetic Stock Center #1P4 |
| **SPP1** | Bacillus phage | Bacillus Genetic Stock Center #1P45 |
| **SP10** | Bacillus phage | The Félix D'Hérelle Reference Center for Bacterial Viruses HER 148 |
| **SP50** | Bacillus phage | The Félix D'Hérelle Reference Center for Bacterial Viruses HER 174 |
| **Goe2** | Bacillus phage | Robert Hertel[1] |

**Supplementary Table S2. Chemical used in this study**

| Chemicals | Source | Identifier |
|---|---|---|
| **LB broth** | VWR | Cat#90003-350 |
| **Dehydrated Agar** | Fisher | Cat#DF0140-07-4 |
| **$MnCl_2$** | Sigma Aldrich | Cat#M3634 |
| **$MgCl_2$** | Sigma Aldrich | Cat#63068 |
| **$CaCl_2$** | Sigma Aldrich | Cat#C7902 |
| **Yeast extract** | Fisher | Cat#DF0127-07-1 |
| **Malt extract** | Fisher | Cat#DF0186-17-7 |
| **Dextrose** | Sigma Aldrich | Cat#G7528 |
| **$FeCl_3$** | Sigma Aldrich | Cat#236489 |
| **$FeSO_4$** | Fisher | Cat#AC201392500 |
| **Ammonium formate** | Sigma Aldrich | Cat#156264 |
| **Formic acid (ACS)** | Sigma Aldrich | Cat#F0507 |
| **Ammonium hydroxide** | | |
| **Formic acid (HPLC)** | VWR | Cat#PI85178 |
| **NH4HC$O_3$ (HPLC)** | VWR | Cat#BJ40867 |
| **Ferrichrome** | Sigma | Cat#F8014 |
| **Linear enterobactin** | Purified from *E. coli*[2] | |
| **Enterobactin** | Gift from Giedroc Lab (Indiana University), synthesized by Nolan Lab (MIT) | |
| **EDDHA** | Sigma Aldrich | AMBH2D6F6CF9 |
| **Erythromycin** | Neta Scientific | Cat# PHR1039 |
| **Kanamycin** | Sigma Aldrich | Cat# K1377 |
| **$Ga_2(SO)_4$** | Sigma Aldrich | Cat#254207 |
| **Sodium citrate** | Sigma Aldrich | Cat# C8532 |
| **Ferrioxamine E** | Sigma Aldrich | Cat#38266 |
| **Nalidixic acid** | Sigma Aldrich | Cat#N8878 |

**Supplementary Table S3. Primers used in this study**

| Primer | Description | Sequence |
|---|---|---|
| **16S_F** | Forward primer for 16 S rRNA | AGAGTTTGATCCTGGCTCAG |
| **16S_R** | Reverse primer for 16 S rRNA | ACGGCTACCTTGTTACGACTT |
| **KanR774** | Forward primer for Kan$^R$ mutant | AGTAAGTGGCTTTATTGATCTTGGG |
| **ErmR815** | Forward primer for Erm$^R$ mutant | CCTTAAAACATGCAGGAATTGACG |
| **spo0A_3pR** | Reverse primer for Δ*spo0A* mutant | TCGAAATTCACGGTGCAAAC |
| **spo0B_3pR** | Reverse primer for Δ*spo0B* mutant | GATTGGCAACCTCAAACAGAA |
| **spo0F_3pR** | Reverse primer for Δ*spo0F* mutant | GGTTTGAAGAACCAAATTCACG |
| **spoIIE_3pR** | Reverse primer for Δ*spoIIE* mutant | GCACTGGGAATATAGGCATCATC |
| **spoIIAA_3pR** | Reverse primer for Δ*spo0F* mutant | AAATCGCTGATCGCTTCTTTC |
| **sigF_3pR** | Reverse primer for Δ*spoIIE* mutant | CTTGAAACCTCCTTATGAATGGTC |

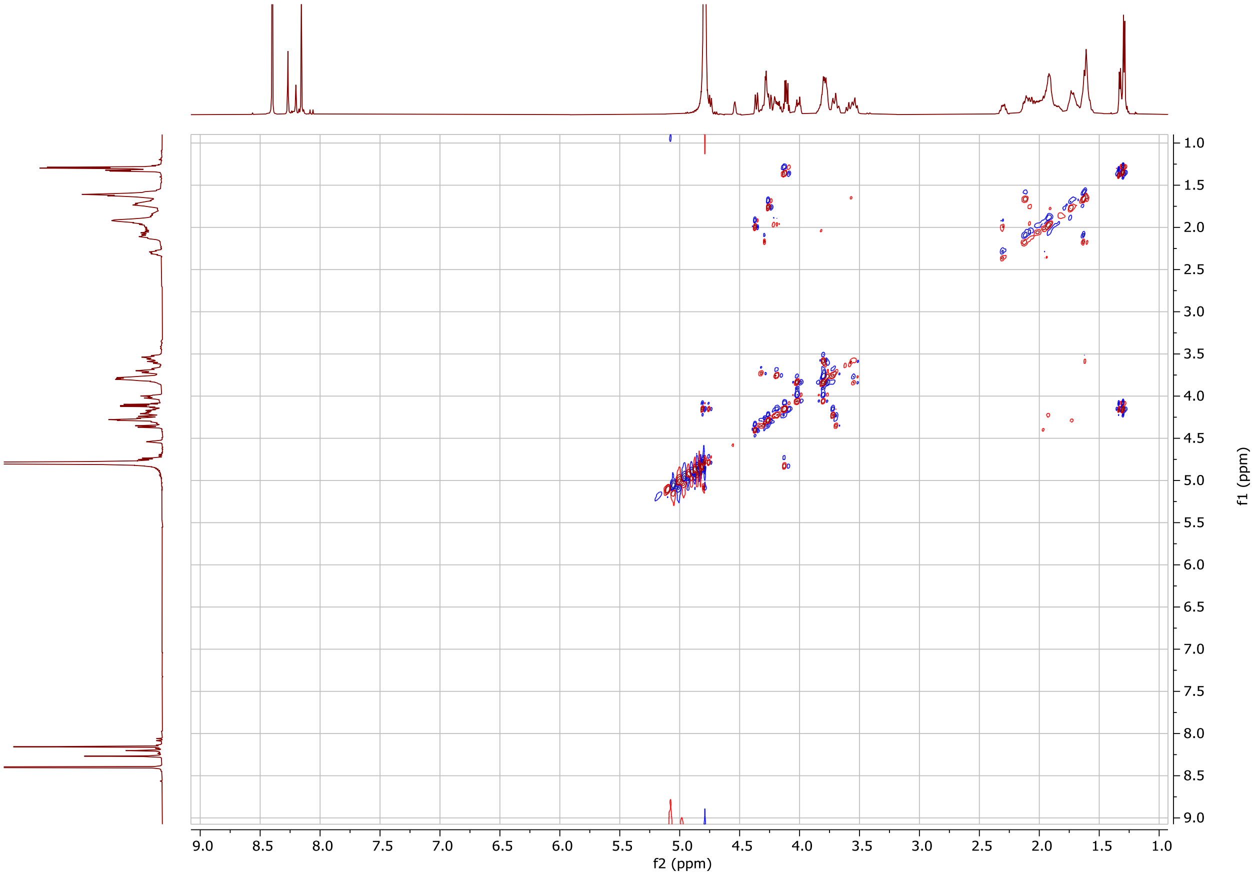

31 **Supplementary Figure S1. DQF-COSY NMR spectrum of Ga-coelichelin in $D_2O$ (500**
32 **MHz).**
33

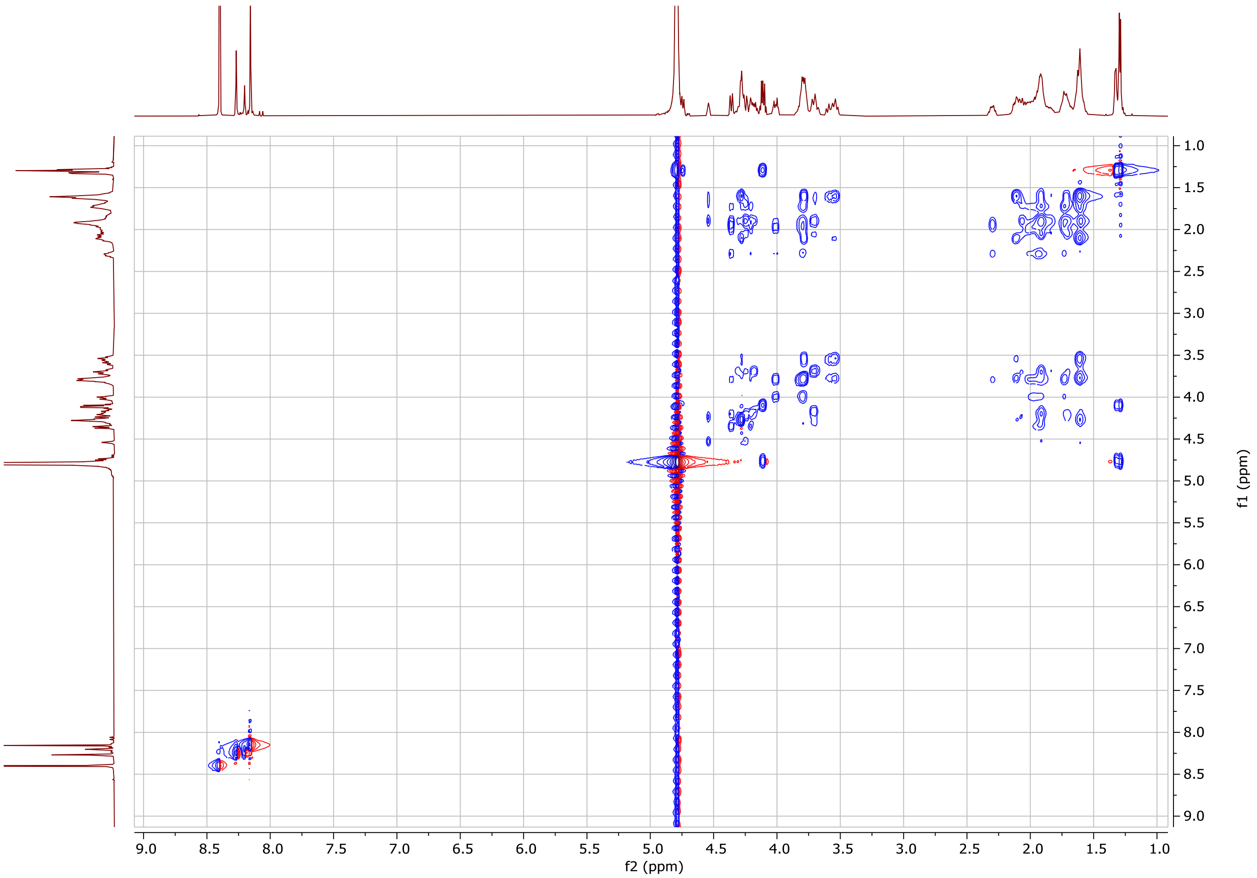

**Supplementary Figure S2. TOCSY (mixing time of 60 ms) NMR spectrum of Ga-coelichelin in $D_2O$ (600 MHz).**

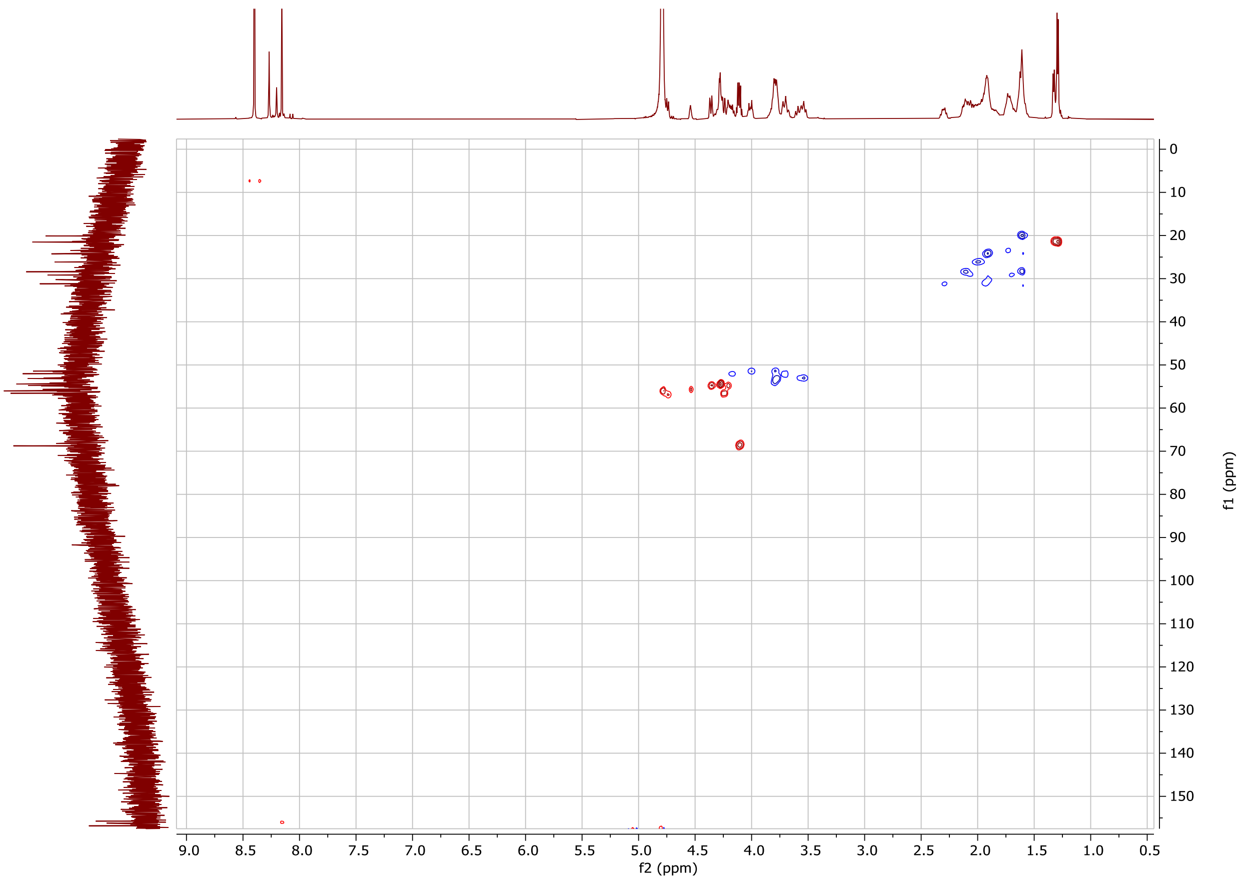

**Supplementary Figure S3. HSQC NMR spectrum of Ga-coelichelin in $D_2O$.**

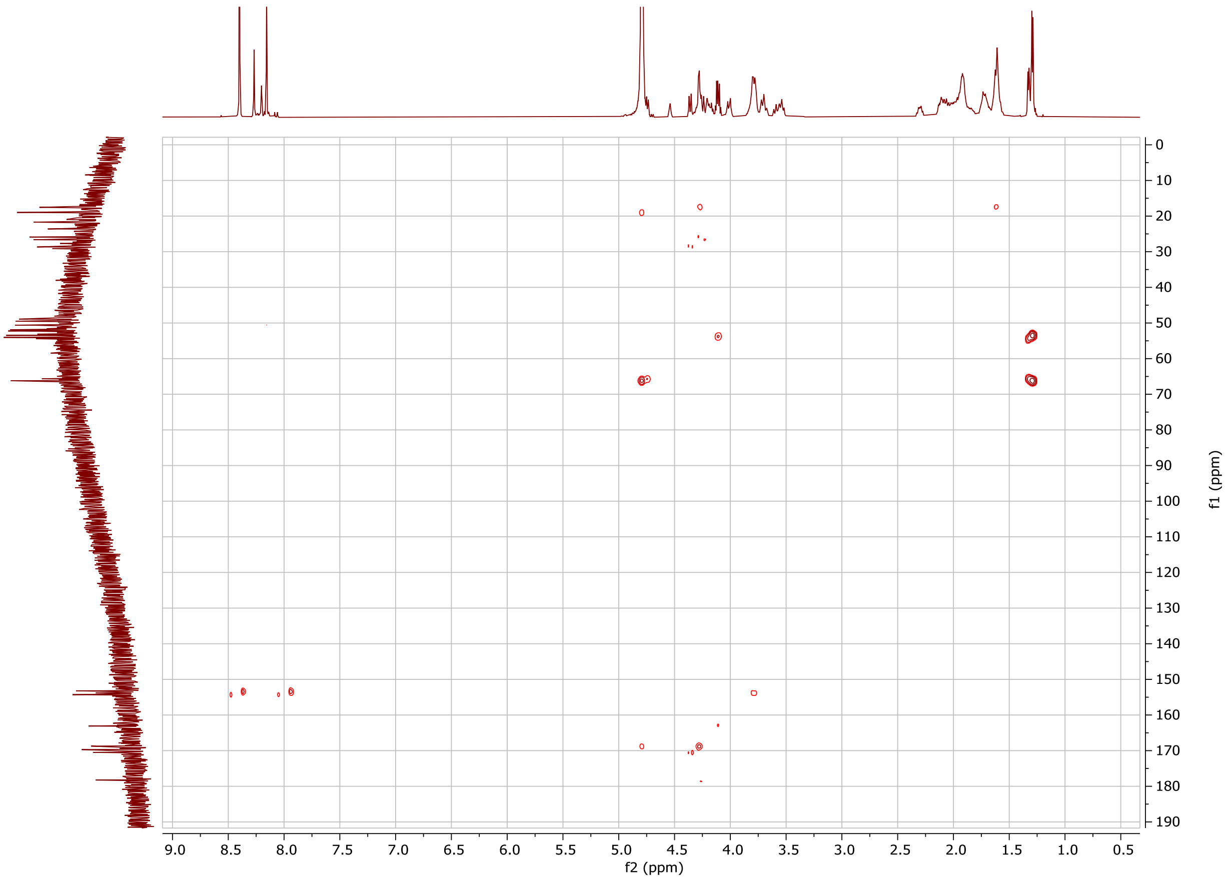

**Supplementary Figure S4. HMBC NMR spectrum of Ga-coelichelin in $D_2O$.**
